# Human cerebrospinal fluid sample preparation and annotation for integrated lipidomics and metabolomics profiling studies

**DOI:** 10.1101/2022.11.07.515425

**Authors:** Kourosh Hooshmand, Jin Xu, Anja Hviid Simonsen, Asger Wretlind, Andressa de Zawadzki, Karolina Sulek, Steen Gregers Hasselbalch, Cristina Legido-Quigley

**Affiliations:** System Medicine, Steno Diabetes Center Copenhagen, Herlev, Denmark; Institute of Pharmaceutical Sciences, Faculty of Life Sciences and Medicine, King’s College London, Franklin-Wilkins Building, 150 Stamford Street, London SE1 9NH, UK; Danish Dementia Research Centre, Copenhagen University Hospital, Rigshospitalet, Copenhagen, Denmark

**Author notes:** Corresponding Author. Steno Diabetes Center Copenhagen, Borgmester Ib Juuls Vej 83, 2730 Herlev, Denmark. (C. Legido-Quigley).

## Abstract

**Objective:** Mass spectrometry (MS)-based lipidomics and metabolomics approaches play an essential role in identifying molecular profiles and relevant clinical biomarkers associated with diseases. Cerebrospinal fluid (CSF) is a metabolically diverse biofluid and a key specimen for exploring biochemical changes in neurodegenerative diseases because its composition reflects brain metabolic activity. CSF lipidomics is receiving increasing attention owing to the importance of lipids in brain molecular signaling and their association with several neurological diseases. Detecting lipid species in CSF using MS-based techniques remains challenging because lipids are highly complex in structure and their concentrations span over a broad dynamic range. This work aimed to develop a robust lipidomics and metabolomics method based on commonly used two-phase extraction systems from human CSF samples.

**Methods:** Prioritizing lipid detection, biphasic extraction methods, Folch, Bligh & Dyer (B&D), Matyash and acidified Folch and B&D (aFolch and aB&D), were compared using 150 μl of human CSF samples (n=6) for the simultaneous extraction of lipids and metabolites with a wide range of polarity in a single extraction. Multiple chromatographical separation approaches, including reversed-phase liquid chromatography (RPLC), hydrophilic interaction liquid chromatography (HILIC), and gas chromatography (GC), were utilized to characterize human CSF metabolome through MS-based untargeted approaches.

**Results:** A total of 219 lipids across 12 lipid subclasses were identified in CSF samples using RPLC-MS/MS. The aB&D method was found as the most reproducible technique (RSD <15%) for lipid extraction. We found remarkable differences in extraction efficiencies among the five different procedures. The aB&D and B&D yielded the highest peak intensities for targeted lipid internal standards and displayed superior extracting power for major endogenous lipid classes. A total of 674 unique metabolites with a wide polarity range were annotated in CSF using, combining RPLC-MS/MS (n=219), HILIC-MS/MS (n=304) and GC-QTOF MS (n=151).

**Conclusions:** Overall, our findings show that the aB&D extraction method provided suitable lipid coverage, reproducibility, and extraction efficiency for global lipidomics profiling of human CSF samples. In combination with RPLC-MS/MS lipidomics, complementary screening approaches enabled a comprehensive metabolite signature that can be employed in an array of clinical studies.

**Graphical abstract:** 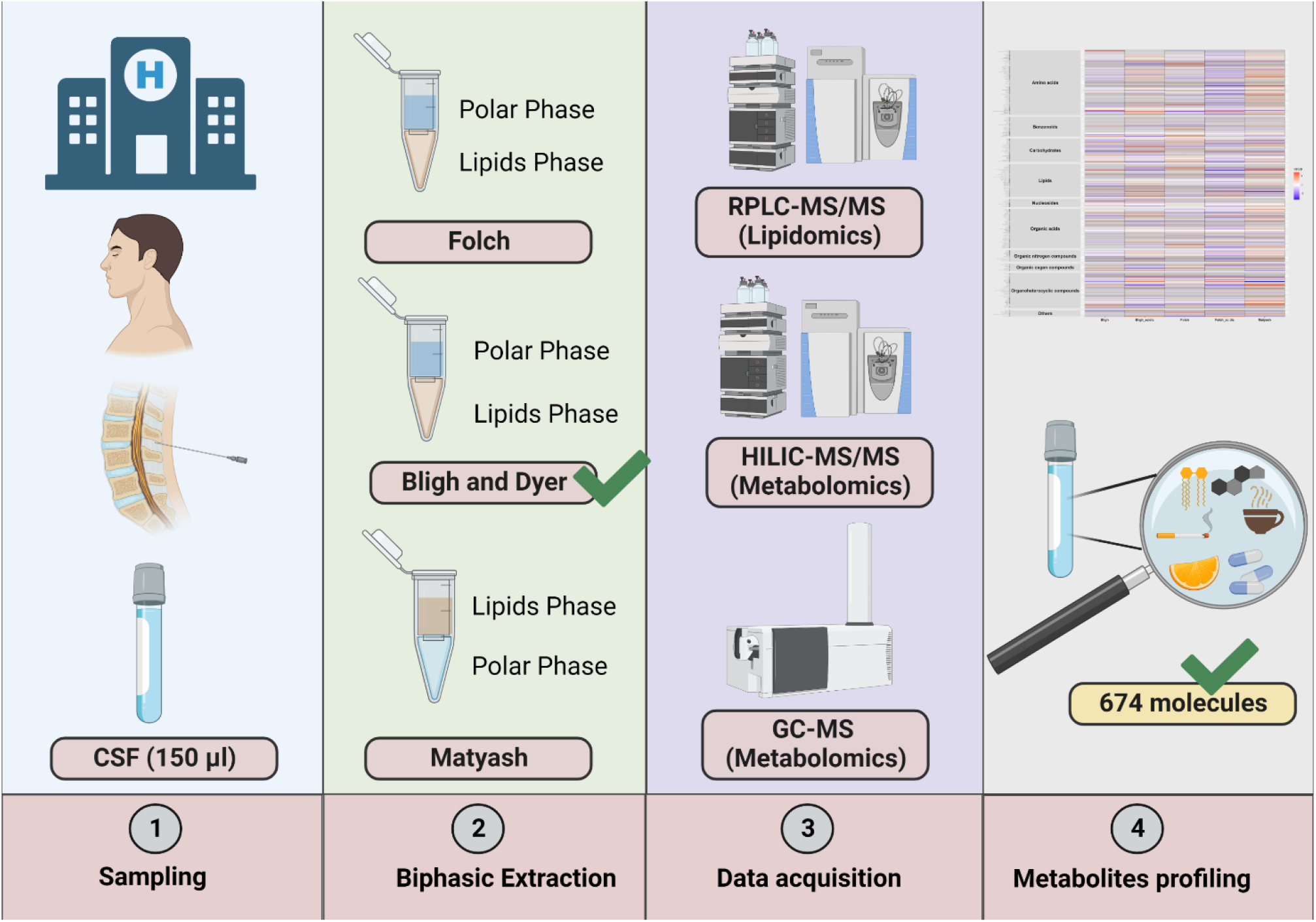

## 1. Introduction

CSF is a clear aqueous fluid that surrounds the brain and spinal cord and fills the ventricular system of the central nervous system (CNS). The CSF serves as a medium for delivering nutrients to neuronal cells, distributing neurotransmitters throughout the CNS, and acts as a lymphatic system to eliminate waste products of cellular metabolism. Due to the direct contact with the brain, CSF is regarded as an important diagnostic tool for identifying novel biomarkers for the diagnosis or prognosis of neurodegenerative diseases.

Lipids are important structural components of cell membranes, serving as energy storage, and play essential roles in controlling and regulating cellular and neuronal functions associated with human physical and pathological conditions [1]. Lipids exhibit remarkable structural diversity characterized by acyl chain lengths, degree of saturation, and a variable head group linked to the phosphate group in glycerophospholipids [2–4]. The diversity of lipids is essential to their cellular functions [4]. Lipidomics aims to analyze, comprehensively and quantitatively, wide arrays of lipids in biological samples [4]. Among the techniques for screening brain-related biomolecules in CSF, lipidomics is receiving increasing attention due to the importance of lipids in brain molecular signaling and their association with several neurological diseases [5–9].

The total lipid content of human CSF is estimated to be 0.2 % of serum levels [10, 11], necessitating the employment of highly sensitive and specific methods for the analysis of lipid species in CSF. The comprehensive analysis of lipids in biological samples using MS-based techniques poses a constant challenge owing to vast structural diversity, a wide concentration range, different physicochemical properties, and the presence of isobaric and isomeric lipid species.

Sample preparation and lipid extraction with a suitable organic solvent mixture are critical steps in quantitative and qualitative lipidomics analysis. One of the most frequently used methods for lipids extraction from various biological fluids is the two-phase extraction methods reported by Bligh and Dyer (B&D), Folch, and Matyash [12–14]. In these methods, an aqueous solution of biological material (e.g., CSF, plasma, serum, urine) is mixed with non-polar (chloroform or MTBE) and polar (methanol and water) solvents, yielding two immiscible-phase systems. This method results in the fractionation of the metabolites into lipid-rich organic and polar aqueous fractions, which can be analyzed separately to increase analyte coverage in a single extraction, thus, opening the way for a more complete characterization of human diseases.

Even though the conventional biphasic extraction methods are suitable for extracting major membrane phospholipids, it is very poor in extracting hydrophilic lipids such as lysophospholipids. Acidification of the extraction solvent can enhance extraction efficiency by neutralizing the negatively charged phosphate group on the phospholipids and causing them to remain in the organic phase [15]. To date, numerous studies have predominantly focused on the comparison of common lipid extraction methods in biofluids such as plasma, serum, and urine [16, 17]. However, a relatively limited number of studies have investigated the effectiveness of various existing biphasic extraction protocols and acidification on lipids and polar metabolites profiling of human CSF using untargeted MS-based lipidomics and metabolomics approaches [16].

Here using an untargeted lipidomics approach on human CSF samples, we find that the most efficient lipids extraction method using two-phase extraction was aB&D. In addition, we followed up with complementing the lipidome with metabolome coverage using RPLC-MS/MS, HILIC-MS/MS, and GC-QTOF MS platforms.

## 2. Materials and methods

### 2.1. CSF samples collection and storage

CSF samples from participants were collected by lumbar puncture as part of their diagnostic evaluation at the Memory Clinic (Copenhagen University Hospital, Rigshospitalet, Denmark). Anonymized leftover samples (n=6) not used for routine analyses were centrifuged (10 min, 2000g, +4°C) and stored in polypropylene tubes at −80°C until analysis.

### 2.2. Chemicals and reagents

Refer to Supplementary Information for a detailed account of chemicals and reagents.

### 2.3. Sample extraction for lipidomics and polar/semi-polar metabolites

Solutions containing lipid internal standards (ISTDs)(PC 34:0, PE 34:0, 17:0 SM (d18:1/17:0), C17 Cer (d18:1/17:0), LPC 34:0, TG 45:0, DG 30:0, PC (14:0/d13), PG 34:0, and PS 34:0) were prepared as a mixture of concentration of 10 ppm (μg/ml) in chloroform: methanol (2:1, v/v). For all experiments, the mixture of internal standards was spiked into the samples before extraction.

Lipids ISTDs mixture solutions were added to an empty Eppendorf tube with the purpose of obtaining the concentration level of 500 ng/mL in the final extract. The tubes were evaporated to dryness prior to extraction using a Biotage TurboVap LV (Uppsala, Sweden) supplied with lab nitrogen. Extractions were conducted in six replicates (150 μL CSF from 6 individual subjects, n=6). The CSF from the same individual was used for comparing different extraction methods. Method blank samples were created using 150 μL of LC-MS grade water instead of CSF. Samples were separated into hydrophobic and hydrophilic portions using a biphasic extraction method. Each extraction procedure was performed as follows:

### 2.4. Extraction according to Folch and acidified Folch (aFolch) method

150 μL CSF was transferred to 2ml Eppendorf tubes with dried ISTDs and mixed with a 1000 μL Folch mixture (CHCl3/MeOH 2:1 v/v). Tubes were vortexed for 10 sec. Phase separation was induced by adding 200 μL of water. The mixture was shaken for 20 min at 1000 RPM at 4°C and centrifugated at 10000 g, for 5 min, at 4 °C to remove any precipitate from the supernatant. From the lower organic phase, 400 μL was collected into a clean microcentrifuge tube for lipidomics analysis. From the upper aqueous phase, 200 μL was collected in a fresh microcentrifuge tube for analysis of hydrophilic metabolites (HILIC method). A portion from the remaining upper (200 μL) and lower layers (200 μL) were taken and combined for metabolomics analysis by GC-QTOF MS. All fractions were evaporated to dryness using a nitrogen evaporator under a gentle stream of nitrogen at room temperature and stored at −80 °C until further analysis. Modification of the Folch method was performed by acidifying the extraction solvent with 1% acetic acid.

### 2.5. Extraction according to B&D and acidified B&D (aB&D) method

In 2ml Eppendorf tubes with dried ISTDs, 150 μL CSF was transferred, and mixed with 900 μl (CHCl3/MeOH 1:2 v/v). The tubes were vortexed for 10 sec, and subsequently, 300μL chloroform was added. The tubes were mixed thoroughly for 10 sec, followed by the addition of 300 μL of water to induce the phase separation. The tubes were shaken for 20 min at 1000 RPM at 4°C and centrifugated at 10000 g, for 5 min, at 4 °C to remove any precipitate from the supernatant. From the lower organic phase, 400 μL were collected into a clean microcentrifuge tube for lipidomics analysis. From the upper aqueous phase, 200 μL was collected into a fresh microcentrifuge tube for analysis of hydrophilic metabolites (HILIC method). A portion from the remaining upper (200 μL) and lower layers (200 μL) were combined for metabolomics analysis by GC-QTOF MS. All fractions were evaporated to dryness using a nitrogen evaporator under a gentle stream of nitrogen at room temperature and stored at −80°C until further analysis. Modification of the B&D method was performed by acidification of the extraction solvent with 1% acetic acid.

### 2.6. Extraction with methyl tertiary butyl ether (MTBE, Matyash method)

In 2ml Eppendorf tubes with dried ISTDs, 150 μL CSF was transferred and mixed with 225 μl cold methanol. The tubes were vortexed (10 sec), followed by the addition of 750 μL MTBE. The tubes were shaken for 20 min at 1000 RPM at 4°C and 188 μL water was added. All the tubes were vortexed and centrifuged at 10000 g, for 5 min, at 4 °C to induce the phase separation. From the upper organic phase, 400 μL was collected into a clean microcentrifuge tube for lipidomics analysis. From the lower aqueous layer, 200 μL was collected into a fresh microcentrifuge tube for analysis of hydrophilic metabolites (HILIC method). A portion from the remaining upper (200 μL) and lower layers (200 μL) were combined for metabolomics analysis by GC-QTOF MS. All fractions were evaporated to dryness using a nitrogen evaporator under a gentle stream of nitrogen at room temperature and stored at −80 °C until further analysis.

### 2.7. Chromatographic and mass spectrometric conditions for lipidomics analysis

RPLC-MS/MS was used to perform lipidomic analysis. Each dried aliquot of the non-polar layer was reconstituted in 50 μl of methanol: toluene (9:1, v/v) (Fisher Scientific) and transferred to 2 mL LC-MS amber vials with 300ul glass inserts. Samples were analyzed using a Vanquish Ultra high-performance liquid chromatography (UHPLC) system coupled with a Q Exactive Plus Hybrid Quadrupole-Orbitrap mass spectrometer in positive and negative electrospray ionization (ESI) modes (Thermo Fisher Scientific, Bremen, Germany). Both ESI (+) and ESI (−) used the same mobile phase composition of (A) 60:40 v/v acetonitrile: water and (B) 90:10 v/v isopropanol: acetonitrile. For ESI (+), 10mM ammonium formate and 0.1% formic acid (Sigma-Aldrich) were added to both solvents A and B. For ESI (−), the modifier was 10mM ammonium acetate. 2 μL of each extracted sample were injected into a reversed-phase Acquity UPLC charged surface hybrid (CSH) C18 column (100 mm length × 2.1 mm internal diameter; 1.7 μm particles) (Waters, Milford, MA, USA) with an Acquity UPLC CSH C18 pre-column (5 mm × 2.1 mm, 1.7 μm particle size) (Waters, Milford MA). Further details on the stepwise gradient used in the LC analysis and a full description of mass spectrometry analysis can be found in the Supplementary Information.

### 2.8. Chromatographic and mass spectrometric conditions for HILIC analysis

Hydrophilic interaction liquid chromatography-tandem mass spectrometry (HILIC-MS/MS) was used to perform HILIC analysis. The dried polar phase was resuspended in acetonitrile: water (4:1, v/v) mixture (50 μl) containing the following internal standards: Glutamine-d5, glutamic acid-d5, phenylalanine-d5, tryptophan-d8, citrulline-d4, alanine-d4, homocitrulline-d3, and leucine-d10 and transferred to 2 mL LC-MS amber vials with glass insert. Samples were analyzed using a Vanquish UHPLC system coupled with a Q Exactive Plus Hybrid Quadrupole-Orbitrap mass spectrometer. Both ESI (+) and ESI (−) used the mobile phase composition of A, 100 % water, and B, 95:5 v/v acetonitrile: water. 10 mM ammonium formate and 0.125% formic acid were added to both solvents. For chromatographic separation of polar and semi-polar metabolites, the resuspended samples were injected (1 μl) onto a Waters Acquity UPLC BEH Amide column (150 mm× 2.1 mm; 1.7 μm) with an additional Waters Acquity VanGuard BEH Amide pre-column (5 mm× 2.1 mm; 1.7 μm) held at 45°C. Further details on the stepwise gradient used in the LC analysis and a full description of mass spectrometry analysis can be found in the Supplementary Information.

### 2.9. Sample derivatization for GC-QTOF MS analysis

Aliquots of previously extracted material (combined upper and lower phases) (400 μl) were transferred to glass vials (1.1ml Microliter Crimp Neck Vial ND11, Mikrolab Aarhus A/S, Denmark), evaporated to complete dryness using nitrogen evaporators to remove any residual water, and capped with magnetic crimp cap (Agilent technologies, Germany). Dried samples were methoximated by the addition of methoxyamine hydrochloride (25 μL) and incubated at 45 °C with agitator shaking (60 min), followed by trimethylsilyl derivatization by the addition of MSTFA containing 1% TMCS (25 μL) and incubated at 45 °C with agitator shaking for 60 min. MSTFA contained an alkane retention index standard mixture (C11, C15, C17, C21, C25, C29, and C33). Lastly, 50 μl of an injection standard 4,4′-dibromooctafluorobiphenyl (concentration = 10 mg/l in hexane) was added to the mixture. All samples were analyzed within 24h of derivatization to avoid degradation in the autosampler.

### 2.10. GC-QTOF MS analysis of CSF extracts

Derivatized samples were analyzed on an Agilent 8890 gas chromatography system (Santa Clara, CA, USA) equipped with a Restek Rxi-5-ms column (30-meter length × 0.25-mm internal diameters (id); 0.25 μm film) with an additional 1.5 m integrated FS-deactivated guard column (Restek, Bellefonte PA). Mass spectrometry was performed using an Agilent 7250 QTOF mass spectrometer, which was controlled by Agilent MassHunter Workstation software (GC/MS Data Acquisition Version 10.0). 1 μl of each sample was injected into the inlet (equipped with a glass wool liner) in the split ratio of 1:5. After every 40 injections, the liner was changed to reduce the carry-over background. The chromatographic gradient was run with helium as the carrier gas at a constant flow of 1 ml/min with the following gradient: The oven temperature started at 50 °C (held for 1 min), ramping 20°C/min to 330°C (14 min) and held constant for 5 min [18]. Mass spectrometry data were collected at m/z 50-550 at a scan rate of 20 spectra/sec. The mass spectrometer was operated with a source temperature of 250°C, in electron ionization mode at −70 eV.

### 2.11. LC-MS/MS and GC-MS data processing using MS-DIAL

Data were reported as quantitative ion peak heights. Data processing for data from each of the three analytical platforms (RPLC-MS/MS, HILIC-MS/MS, and GC-MS) was performed using the open-source software MS-DIAL (version 4.8) and included peak detection, deconvolution, feature alignment, gap filling, deisotoping, adduct identification, accurate mass/retention time (m/z-RT) library matching, and MS/MS library matching [19]. Features were annotated using defined confidence levels [20].For lipidomics, experimental MS/MS spectra were matched to reference MS/MS spectra from the LipidBlast library [21]. For HILIC data sets, experimental tandem MS spectra were matched to library spectra from the Mass Bank of North America (MoNA) (http://massbank.us) and NIST 17 MS/MS spectral library to annotate the features [22]. GC-MS spectra were matched against public mass spectral libraries (Fiehn library, In-house library, MassBank Japan, GOLM DB, GNPS, and HMDB). The details of the LC-MS and GC-MS data processing are described in Supplementary Information.

### 2.12. Statistical Analysis

The peak intensity table from the lipidomics and metabolomics analysis was submitted to MetaboAnalyst 5.0 for data normalization [23]. Data not meeting normality assumptions were cube-root-transformed, and Pareto scaled to achieve normal distribution. A supervised partial least square-discriminant analysis (PLS-DA) was used for multivariate statistics and visualization. The significance of differences in the mean value between different extraction methods was assessed by univariate one-way ANOVA analysis and significant differences were compared using Tukey’s HSD using R statistical language (ver. 3.5.3). The packages “ggpubr “ and “tidyverse “ in R were used for data wrangling and visualization [24, 25]. Reproducibility was expressed as the relative standard deviation (%RSD) in replicates (n=6) and was calculated by dividing the standard deviation by the mean of peak intensity and multiplying by 100.

## 3. Results and discussion

### 3.1. Reproducibility and evaluation of extraction procedures

The ability of biphasic extraction methods to extract complex lipids was evaluated using reproducibility and relative abundance of lipids signal. Reproducibility and extraction efficiency of each extraction method were calculated using CSF replicate samples (n = 6) spiked with ten non-endogenous lipid internal standards (their absence in CSF was confirmed by LC-MS) representing the nine main lipid families (PC, LPC, DG, Cer, TG, SM, PS, PG, and PE). Among the five tested extraction techniques, the aB&D method yielded the lowest variations in the peak intensities (RSD% for exogenous lipid species ranging from 5 % to 14 %) (**Figure 1**), demonstrating a high level of reproducibility of this method. Among the lipid families, the highest variations were observed for neutral lipid specie DG with RSD values of 18%. The B&D and aB&D methods yielded better extraction efficiency values for all the lipid classes showing substantially higher peak intensities for the ten targeted lipid species (**Figure 1**) compared to Folch and Matyash methods. Acidification negligibly impacted extraction efficiencies of certain lipid for the B&D method, exhibiting slightly higher peak intensity for PE 34:0 (**Figure 1**). ANOVA pairwise multiple comparisons were used to assess whether a significant difference existed between different lipid extraction methods (**Supplementary Table 1**). No significant difference was found between B&D and aB&D as well as Folch and aFolch for all the lipid species (p >0.05). However, a significant increase in the lipid peak intensities was observed comparing B&D with Folch and Matyash methods, with p <0.05.

**Figure 1.**
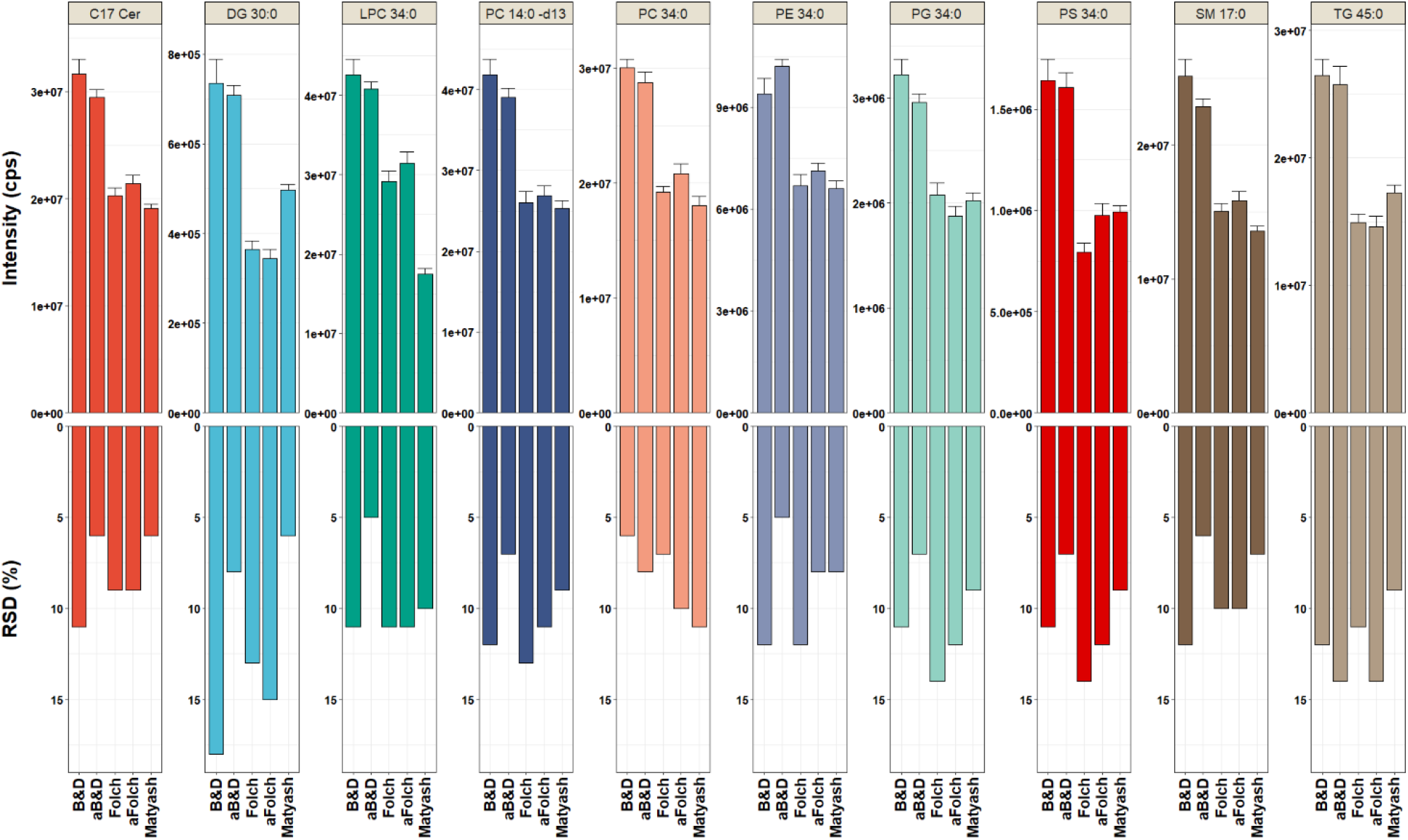
Comparison of non-endogenous lipid species extracted from CSF. A) Comparison of extraction methods using the average peak intensities of targeted lipid internal standards. Y axis indicates the signal intensity. Error bars indicate the standard error of the mean of six independent biological replicates. B) Reproducibility of different extraction methods as measured from relative standard deviation (%RSD) of peak intensities for 10 targeted lipids (n=6). Abbreviations, acidified Bligh & Dyer: aB&D; acidified Folch: aFolch; C17 Cer (ceramide (d18:1/17:0)); DG 30:0 (diacylglycerol 15:0/15:0); LPC 34:0 (lysophosphatidylcholine 17:0/17:0), PC (14:0/d13) (phosphatidylcholine (14:0/d13)), PC 34:0 (phosphatidylcholine 17:0/17:0); PE 34:0 (phosphatidylethanolamine 17:0/17:0); PG 34:0 (phosphatidylglycerol 17:0/17:0); PS 34:0 (lysophosphatidylserine 17:0/17:0); SM 17:0 (sphingomyelin 17:0/17:0); TG 45:0 (triacylglycerol 15:0/15:0/15:0).

The relatively high repeatability was accompanied by high peak intensities of exogenous lipid species for the aB&D method, suggesting that this method is well suited for untargeted lipidomics of human CSF samples. This finding may indicate that with the B&D extraction, targeted lipids partition better into the organic phase. Our finding contrasted with results observed in a previous study reported by Reichl et al., 2020, showing that the aFolch method was the most suitable for the efficient extraction of lipid species [16]. The differences between the two studies might stem from using of different exogenous lipid species as ISTDs. In agreement with the observation made by Reichl et al. (2020), acidification was beneficial for recovering certain lipids.

### 3.2. Lipidomics analysis of CSF and comparing lipid extraction methods

In total, over 32000 features were detected in human CSF samples in positive and negative ionization modes using RPLC-MS/MS. After blank subtraction and removal of isotopes, duplicates, and adducts, 219 distinct features were identified and quantified in both polarities **(Figure 2)**. Among the five biphasic extraction systems examined in this study, 12 lipid classes were detected in CSF. TG was the class with the highest number of lipids (n=67), followed by glycerophosphocholines (n=52), SM (n=22), PE (n=20), glycosphingolipids (GlcCer) (n=13), glycerophosphoinositols (PI) (n=11), lyso-glycerophosphoethanolamines (LPE) (n=7), Cer (n=7), LPC (n=6), cholesteryl esters (CE) (n=5), DG (n=4), sterols (n=3), and monoacylglycerols (MG) 1 species **(Supplementary Table S2)**. Within glycerophosphocholines, PC species (n=34), PC plasmalogens (n=13) and ether-linked PC (n=5) subclass were identified. Within glycerophosphoethanolamine, PE (n=8) and PE plasmalogens (n=12) subclasses were identified. Hence, the family of glycerophospholipids (PC, LPC, PE, LPE, and PI) exhibited the greatest number of species that were collectively detected in CSF. PC 34:1, MG 8:0, PC 32:0, PC 34:2, and SM d34:1 were found to be the five most abundant lipids detected in human CSF, respectively, which was in line with the previous study [26]. The least abundant lipid in CSF was LPE 16:0 and LPE 18:2. The major lipids detected in human CSF are listed in **Supplementary Table S3**.

**Figure 2.**
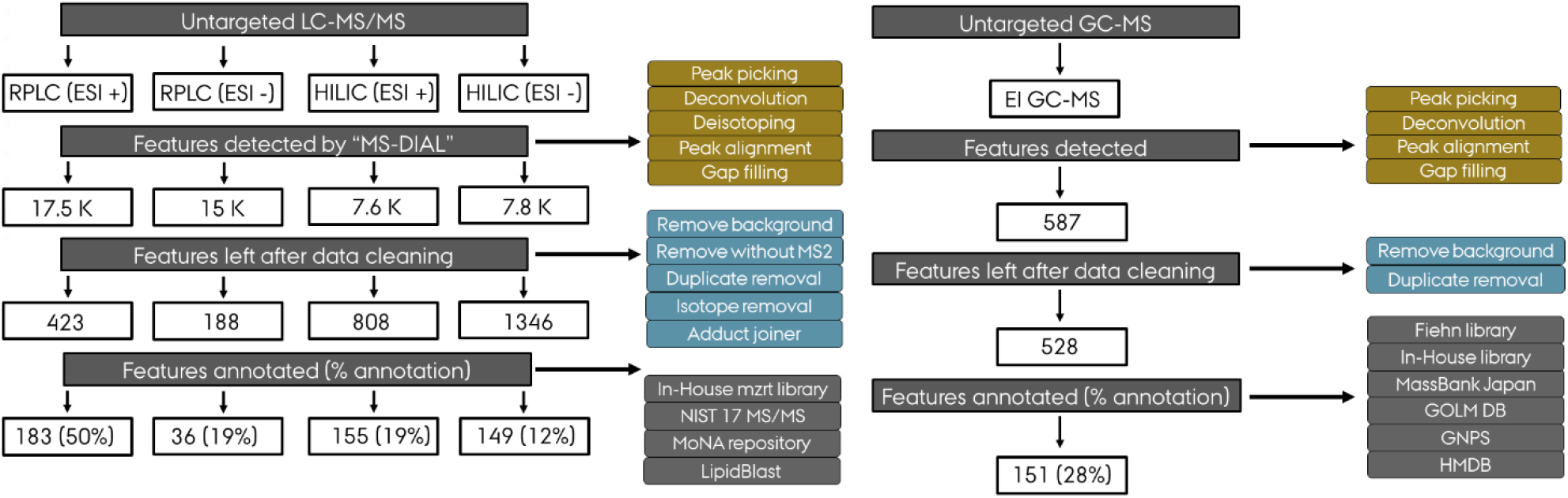
Schematic of the workflow used with MS-DIAL program to process and annotate the metabolites detected by three different analytical platforms (RPLC-MS/MS, HILIC-MS/MS and GC-QTOF MS). Abbreviations, reversed-phase liquid chromatography (RPLC); hydrophilic interaction liquid chromatography (HILIC); and gas chromatography (GC), EI (electron Impact Ionization).

There are several noteworthy reports in which CSF lipids have been successfully detected. Using the Matyash extraction method, Wood et al. (2018) identified 23 lipids in 500 μl of CSF using direct infusion analyses coupled with an orbitrap mass spectrometer [27]. Reichl et al. 2020 used the Folch extraction method and an LC column coupled to ion mobility quadrupole time of flight instrument to obtain untargeted global lipidomics analysis of CSF (75 μl), resulting in the annotation of 38 lipids [16]. In another study, Iriondo et al. 2019 reported that employment of the isopropanol protein precipitation method led to the annotation of 175 different lipid species in 200 μl of CSF utilizing the LC-QTOF system [17]. Lipidomics analysis of CSF using an LC-Q Exactive Mass Spectrometer resulted in the detection of 122 and 133 lipids in 75 and 120 μl of CSF in two studies using aFolch and methanol extraction methods, respectively [9, 26]. Byeon al. (2021), detected a great number of lipid species using 400 μl CSF sample material [28]. The previous work and the quest to detect as many lipids as possible in CSF is very important, as lipids are key mediators found in the pathogenesis of various neurological disorders [5–9].

A heatmap was used to assess the differences between lipid biphasic extraction methods of a non-polar fraction. The heatmap comparing average peak intensities revealed that the levels of endogenous lipid species differed considerably between the different extraction methods (**Figure 3, Supplementary Table S4)**. Using B&D and aB&D method substantially increased the abundance of a broader range of lipid species, including major phospholipids, certain sphingolipids, and highly lipophilic neutral lipids (cholesteryl ester, and triacylglycerols), as compared to Folch and Matyash methods. Thus, this observation suggests that B&D and aB&D methods are suitable for extracting major lipids from CSF. The superiority of the B&D methods for the simultaneous extraction of lipids could be explained by the broad polarity of the extraction solvent [29]. The Folch method was particularly superior for PI (40:4) and PI (40:5) compared to other methods (**Figure 3)**. The Matyash method had relatively high extraction efficiency for certain HexCers. However, the Folch method showed moderate extraction efficiencies, and the Matyash method turned out to be deficient in extracting major lipid classes, compared to the tested alternatives (**Figure 3)**. This is likely driven by the physical-chemical characteristics of MTBE compared to chloroform which makes the B&D and Folch method to be more selective for specific classes of lipids [29].

**Figure 3.**
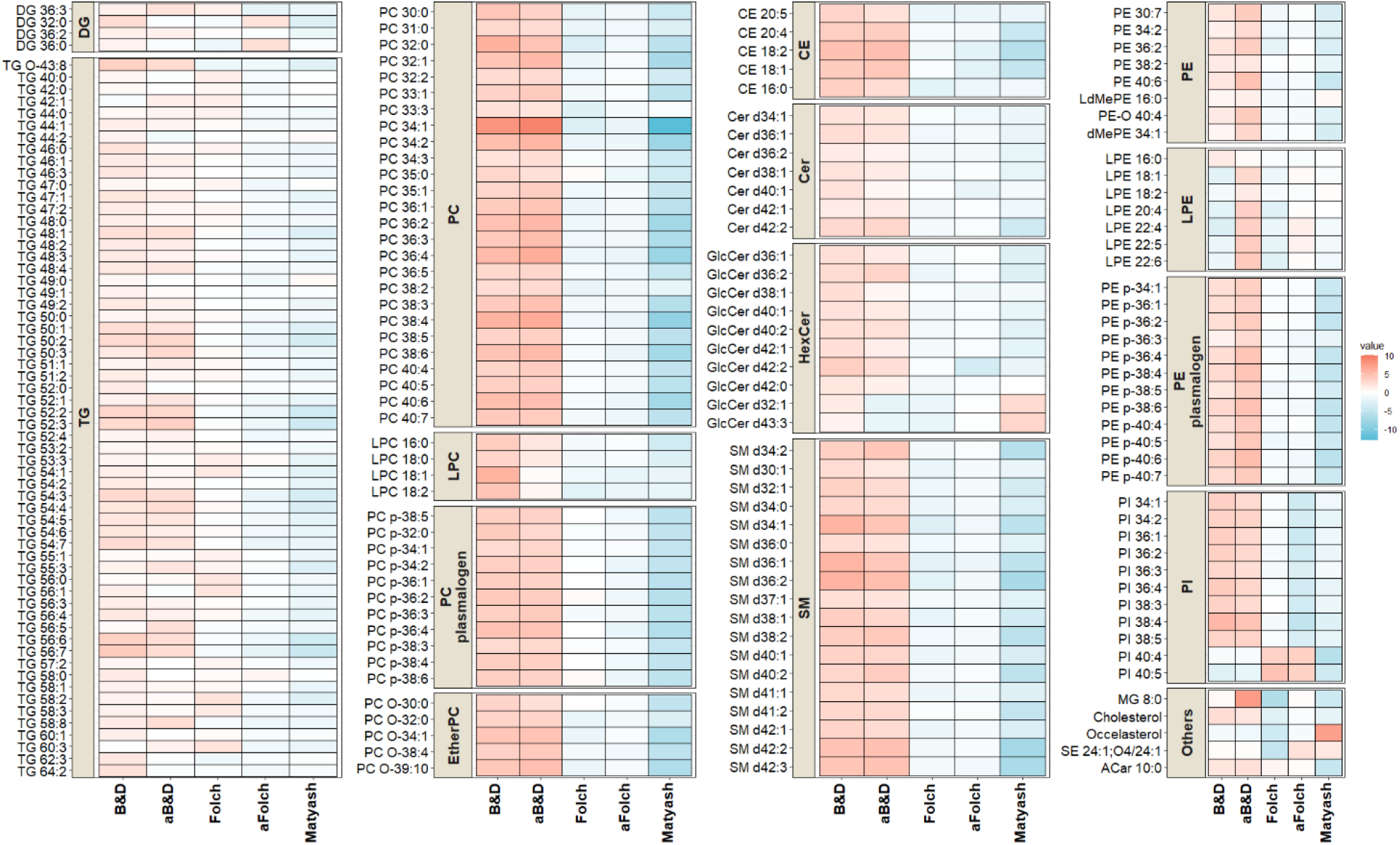
Heatmap comparison of the lipid species among the five different extraction methods. The average intensities of the identified lipids were compared. Red color intensity represents increased values, and blue indicates decreased values. Abbreviations, acidified Bligh & Dyer: aB&D; acidified Folch: aFolch; DG (diacylglycerol); TG (triacylglycerol); PC (phosphatidylcholine); LPC (lysophosphatidylcholine); CE (cholesterol ester); Cer (ceramide); HexCer (hexosylceramide); SM (sphingomyelin); PE (phosphatidylethanolamine); LPE (lyso-phosphatidylethanolamine); PI (Phosphatidylinositol).

Acidification impacted the extraction efficiencies of lipid species in the B&D method, as shown on the PLS-DA plot comparing the two groups (**Supplementary Figure 1A**). The volcano plot comparing B&D and aB&D showed that the level of polar lipid species like LPE was significantly increased after extraction solvent acidified with 1% acetic acid (**Supplementary Figure 1B, Supplementary Table S4**). In contrast, acidification of the B&D method resulted in a substantial decrease in the level of LPC, but still, their intensity was higher than the other investigated methods. Our finding is in agreement with a previous study showing that the peak intensities of LPE 18:1 (d7) were significantly higher in the B&D method with acetic acid compared to non-acidic B&D, while the level of LPC 18:1(d7) was marginally lower when extracted in an acidified condition [16]. Acidification was previously demonstrated to favor the extraction efficiencies of lipid species [30, 31]. The exposure of lipids to strong acids such as hydrochloric acid (HCl) would favor lipids hydrolysis during the extraction, which results in the generation of artifacts. Acidifying solvent mixture systems with a milder organic acid such as acetic acid, in contrast to stronger acid such as HCL, can not only enhance the extraction efficiency but also minimize the hydrolysis of ester and vinyl ether bonds of phospholipids and plasmalogens [15, 30]. Overall, our results showed that B&D and aB&D methods performed equally robust for lipidomics analysis of CSF compared to Folch and Matyash methods. However, aB&D resulted in lower variation and better reproducibility (**Figure 1**), and it is the best choice for an untargeted biphasic extraction of lipids in the human CSF.

### 3.3. CSF metabolomics analysis by HILIC-MS/MS and GC-QTOF

The advantage of using liquid-liquid extractions is the ability to explore both the organic and aqueous layers for increased analyte coverage in untargeted lipidomics and metabolomics studies using minimal sample material [32]. CSF is a metabolically diverse biofluid, and currently, there is no single analytical platform capable of separating, characterizing, and quantifying all analytes present in a CSF. Hence, a combination of analytical techniques was used to maximize the lipidome and metabolome coverage in the single sample.

For HILIC-MS/MS analysis, out of 15400 detected ions, 2154 chromatographic features with associated MS/MS spectra were detected in both polarities, and 304 of these features were annotated (**Figure 2**). The structurally annotated or identified compounds were from different structural classes such as amino acids and derivatives (n=79), organic acids (n=48), organoheterocyclic compounds (n=43), lipids (n=40), carbohydrates (n=29), benzenoids (n=24), organic nitrogen compounds (n= 14), nucleosides (n=10), and organic oxygen compounds (n=9) **(Figure 4 and Supplementary Table S2)**. The 9 most abundant metabolites found in CSF using HILIC-MS/MS were lactic acid, creatine phosphate, N-acetyl-glutamine, 3-hydroxybutyric acid, glutamine, 3-hydroxyisovaleric acid, myo-inositol, choline, and creatine, respectively. The use of the HILIC-MS technique allowed the identification of 40 unique lipids (with only one overlapped with GC-MS features), including medium-chain fatty acids (n=10), hydroxy fatty acids (n=10), acylcarnitines (n=9), long-chain fatty acids (n=1), hydroxy bile acids (n=1), and cholesterol derivatives (n=1) **(Supplementary Table S2**).

**Figure 4.**
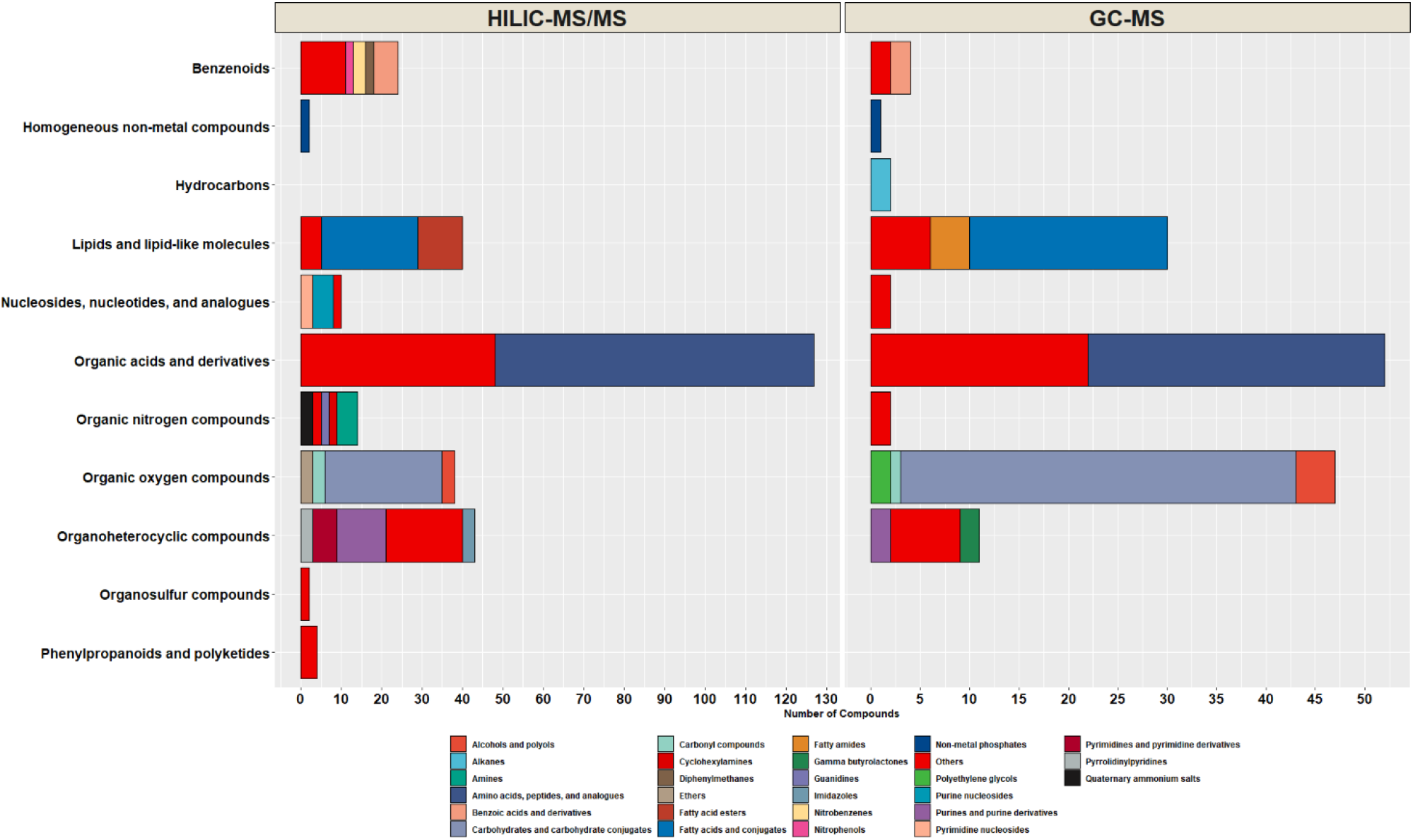
Stacked bar plot showing the chemical classification of the compounds annotated by HILIC-MS/MS and GC-QTOF MS platforms. Class and subclass level ontology is determined by ClassyFire software. The colors in the stack bar plot designate the number of metabolites in each chemical subclass group.

In addition to the detection of endogenous metabolites in human CSF, HILIC-MS/MS allowed the identification and quantitation of unique exogenous compounds, including drugs, vitamins (vitamin B3, B5, B6, and C), and metabolites associated with smoking (nicotine, cotinine, and 3-hydroxycotinine), and coffee (caffeine, and metabolite of caffeine, paraxanthine). The drugs that were found included metformin (medication for the treatment of type 2 diabetes), tramadol (narcotic analgesic), bicalutamide (a drug which is used for the treatment of advanced prostatic cancer), acetaminophen and its metabolization products such acetaminophen sulfate, and 3-hydroxyacetaminophen. We used the Agilent 1290 Infinity UHPLC system coupled to the Agilent 6460 triple quadrupole mass spectrometer to confirm the existence (matching retention time and the intensity ratio of the two MRM transitions) and determine the absolute concentration of some of the exogenous metabolites detected in CSF using aB&D extraction method. The concentration of metformin and bicalutamide in CSF was calculated as 115 ng/ml and 13.5 ng/ml in two independent samples. In addition, in one of the participants, CSF concentrations of nicotine and the main nicotine metabolite, cotinine, were found to be 0.1 and 0.8 ng/ml, respectively.

The presence of exogenous compounds highlights the potential of metabolomics in aiding the stratification of subjects based on their metabolic profiles (e.g., smoker vs. nonsmoker). Once the subjects are successfully stratified into groups, a better understanding of the pathophysiology of their disease can be achieved [33]. In addition, diseases that tend to occur in association with other diseases can be identified (e.g., having diabetes and concurrently developing Alzheimer’s disease). Thus, identifying potential confounding variables that can be risk factors for a disease can provide essential information in characterizing diseases and developing novel targeted therapies and interventions within the context of precision medicine.

GC-TOF-MS analysis yielded a total of 151 identified metabolites (with a wide range of polarity) in human CSF using a aB&D biphasic extraction method. The identified compounds were mainly primary metabolites, including carbohydrates (n=40), amino acids and derivatives (n=30), lipids (n=30), organic acids (n=22), organoheterocyclic compounds (n=11), and organic oxygen (n=7) **(Figure 4 and Supplementary Table S2)**. The 10 most abundant metabolites found in CSF using GC-MS were 2-hydroxypropanoic acid, lactic acid, 4-hydroxybutanoic acid, glucose, iditol, glycerol, citric acid, galacturonic acid, sorbose, and urea **(Supplementary Table S3)**. GC-MS platform allowed the identification and quantification of 30 unique lipids, including medium-chain fatty acids (n=3), long-chain fatty acids (n=10), primary fatty acid amides (n=4), methyl-branched fatty acid (n=3), hydroxy fatty acids (n=3), and cholesterol. When lipidomics and metabolomics results were combined, the three platforms allowed even greater coverage of the human CSF lipids (lipidomics (n=219), HILIC-MS (n=40), and GC-MS (n=30). Among 289 identified lipids, 2 of them were shared between 3 platforms. This result reflects the advantage of performing multi-omics analyses using biphasic extraction for achieving enhanced lipids coverage in untargeted lipidomics and metabolomics studies.

The Venn diagram displays the distribution of features detected by global lipidomics and metabolomics analysis (**Figure 5**). Out of 674 metabolites detected in human CSF samples, GC–MS and HILIC-MS/MS methods identified a common set of 31 compounds. Lipidomics and GC–MS methods identified a common set of 1 compound (cholesterol). Lipidomics detected 218 unique compounds that HILIC-MS/MS and GC–MS methods cannot detect. HILIC-MS/MS detected 273 compounds that GC–MS methods cannot detect. Furthermore, GC–MS detects 119 compounds that HILIC-MS/MS cannot detect. Thus, using a combination of analytical platforms allowed the identification and quantification of 642 unique metabolites. Previous studies demonstrated that using a combination of LC-MS, GC-MS, Fourier transform-MS, nuclear magnetic resonance (NMR), and inductively coupled plasma-(ICP)-MS resulted in the identification of 308 and 476 detectable metabolites in human CSF [34, 35]. Our pipeline yielded broader coverage of the CSF metabolome, reflecting an improvement in the number of identified metabolites in CSF.

**Figure 5.**
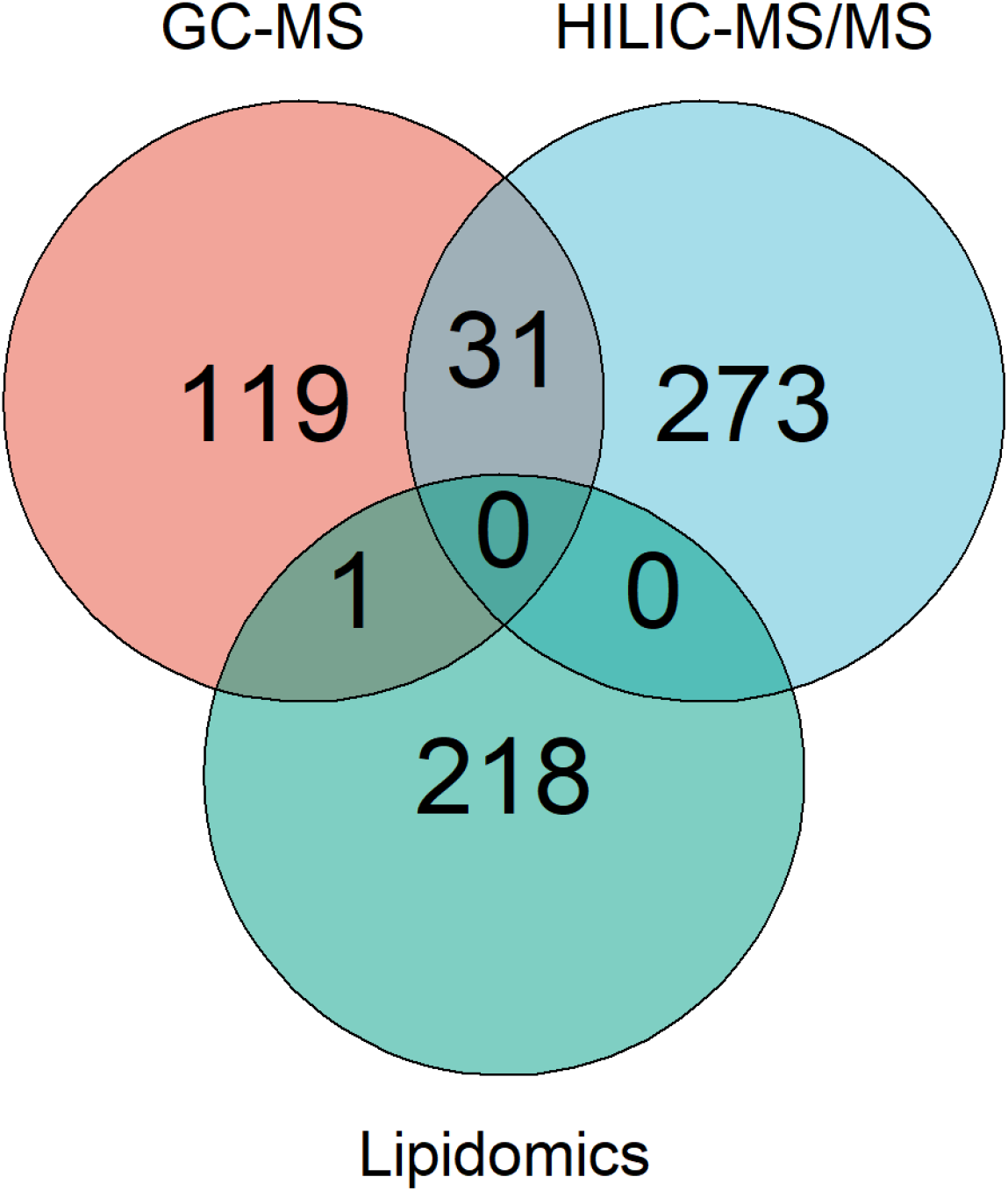
Venn diagram showing the overlap of CSF metabolites detected by RPLC-MS (lipidomics), HILIC-MS/MS (metabolomics), and GC-MS (metabolomics). Abbreviations, reversed-phase liquid chromatography (RPLC); hydrophilic interaction liquid chromatography (HILIC); and gas chromatography (GC), EI (electron Impact Ionization).

Overall, our results demonstrate the strength of optimizing a lipidomics platform and follow up with the detection of a broad range of CSF metabolome with a wider range of polarity. CSF is a metabolically diverse biofluid and can be a rich source of disease targets and biomarkers associated with human disease.

## 4. Conclusions

Here, we show a detailed comparison of frequently used biphasic extraction methods using human CSF samples to understand how the different solvent systems influence the extraction efficiency of lipids. Based on extraction efficiency, reproducibility, and coverage of different lipid classes, Bligh & Dyer method was deemed the best choice for untargeted biphasic extraction for lipidomics in CSF. Moreover, the acidified Bligh & Dyer method has shown good performance for global metabolic profiling methods using three different analytical platforms: RPLC-MS/MS, HILIC-MS/MS, and GC-QTOF MS. Our results also revealed that global lipidomics and metabolic profiling methods could detect 674 distinct metabolites with a wide range of polarities in human CSF from a minimal amount of sample.

## Contributions

KH, CLQ, JX and AHS conceived and planned the research and experiments. KH carried out the study and performed the experiments. KH, KS and AdZ performed the LC-MS and GC-MS analysis. KH, AW and JX analyzed the data and performed statistical analyses. KH and CLQ wrote the manuscript. All authors provided critical feedback and helped to revise the manuscript. All authors have reviewed the manuscript.

## Acknowledgements

The authors thank participants in this study. We would also like to thank Ismo Matias Mattila, and Annette F Bjerre, for their assistance in the laboratory.

