## Supplementary Information for "Human cerebrospinal fluid sample preparation and annotation for integrated lipidomics and metabolomics profiling studies"

### 2.2. Chemicals and reagents

Chemicals and reagents were obtained from the following commercial sources (purity): Lipid internal standards including phosphatidylcholine (PC) 17:0/17:0 (99%), phosphatidylethanolamine (PE) 17:0/17:0 (99%) , lysophosphatidylserine (PS) 17:0 (99%), phosphatidylglycerol (PG ) 17:0 (99%), sphingomyelin (SM) 17:0 SM (d18:1/17:0) (99%), ceramide (Cer) C17 Cer (d18:1/17:0) (99%), lyso-phosphatidylcholine (LPC) 17:0/17:0 (99%), and were purchased from Avanti Polar Lipids, Inc. (Alabaster, Alabama, USA). Triacylglycerol (TG) 15:0/15:0/15:0 (98%), Diacylglycerol (DG) 15:0/15:0 (98%), were purchased from Cayman Chemical (BioNordika A/S, Herlev, Denmark). PC (14:0/d13)) (99%) was purchased from Sigma-Aldrich (Copenhagen, Denmark). Metformin hydrochloride (98%) was obtained from Fisher Scientific (Roskilde, Denmark). Bicalutamide (98%), cotinine (98%), and nicotine (99%) were purchased from Sigma-Aldrich (Copenhagen, Denmark). Glutamine-d5, glutamic acid-d5, phenylalanine-d5, tryptophan-d8, citrulline-d4, alanine-d4, homocitrulline-d3, and leucine-d10 were purchased from CDN Isotopes (Pointe-Claire, Quebec, Canada). Acetonitrile (ACN), isopropanol (IPA), methanol (MeOH), and water (liquid chromatography–mass spectrometry (LC-MS)- grade solvents), formic acid, acetic acid, ammonium acetate, ammonium formate, and methoxyamine (MOX) reagent were obtained from Fisher Scientific (Roskilde, Denmark). LC-MS and GC-MS grade solvents including tert-Butyl methyl ether (MTBE), chloroform ($CHCl_3$), dichloromethane, pyridine, hexane, N-methyl-N-(trimethylsilyl) trifluoroacetamide (MSTFA), and trimethylsilyl chloride (TMCS) were acquired from Sigma-Aldrich (Copenhagen, Denmark).

### 2.7. Chromatographic and mass spectrometric conditions for lipidomics analysis

RPLC-MS/MS was used to perform lipidomic analysis. Each dried aliquot of the non-polar layer was reconstituted in 50 µl of methanol: toluene (9:1, v/v) (Fisher Scientific) and transferred to 2 mL LC-MS amber vials with 300ul glass inserts. Samples were analyzed using a Vanquish Ultra high-performance liquid chromatography (UHPLC) system coupled with a Q Exactive Plus Hybrid Quadrupole-Orbitrap mass spectrometer in positive and negative electrospray ionization (ESI) modes (Thermo Fisher Scientific, Bremen, Germany). Both ESI (+) and ESI (-) used the same mobile phase composition of (A) 60:40 v/v acetonitrile: water and (B) 90:10 v/v isopropanol: acetonitrile. For ESI (+), 10mM ammonium formate and 0.1% formic acid (Sigma-Aldrich) were added to both solvents A and B. For ESI (-), the modifier was 10mM ammonium acetate. 2 µL of each extracted sample were injected into a reversed-phase Acquity UPLC charged surface hybrid (CSH) C18 column (100 mm length x 2.1 mm internal diameter; 1.7 µm particles) (Waters, Milford, MA, USA) with an Acquity UPLC CSH C18 pre-column (5 mm × 2.1 mm, 1.7 µm particle size) (Waters, Milford MA). The column temperature was maintained at 65°C throughout the run with a flow rate of 0.6 mL/min. The linear gradient method was programmed as follows: from 0 min 15% (B), 0–2 min (B) was changed linearly from 15% to 30%, 2–2.5 min (B) changed linearly to 48%, 2.5–11 min (B) increased to 82%, 11–11.5min (B) increased to 99%, 11.5–12min, hold at 99% (B), 12–12.1 min returned to 15% (B), and 12.1–15 min held at 15% (B). The autosampler was kept at 4 °C, and needles were washed with isopropanol for 20 seconds before and after each sample injection. Q-Exactive Plus Hybrid Quadrupole-Orbitrap Mass Spectrometer used the following parameters: The ion spray voltage floating: ±3 kV; sheath gas (nitrogen) flow at 60 units; auxiliary gas flow at 25 units; the capillary temperature at 300°C; auxiliary gas heater temperature at 350°C; and sweep gas flow at 2 units. In addition, full scan MS1 and data-dependent acquisition (DDA) MS/MS were acquired in centroid mode with the following parameter settings: mass range, m/z 65–975, MS1 resolving power, 30,000 full width at half maximum (FWHM), MS2 resolving power, 15,000 FWHM, with the top five ions from each MS1 scan being selected for MS/MS fragmentation by using normalized collision energy at 15, 30, and 45 eV.

### 2.8. Chromatographic and mass spectrometric conditions for HILIC analysis

Hydrophilic interaction liquid chromatography-tandem mass spectrometry (HILIC-MS/MS) was used to perform HILIC analysis. The dried polar phase was resuspended in acetonitrile: water (4:1, v/v) mixture (50 µl) containing the following internal standards: Glutamine-d5, glutamic acid-d5, phenylalanine-d5, tryptophan-d8, citrulline-d4, alanine-d4, homocitrulline-d3, and leucine-d10 and transferred to 2 mL LC-MS amber vials with glass insert. Samples were analyzed using a Vanquish UHPLC system coupled with a Q Exactive Plus Hybrid Quadrupole-Orbitrap mass spectrometer. Both ESI (+) and ESI (-) used the mobile phase composition of A, 100 % water, and B, 95:5 v/v acetonitrile: water. 10 mM ammonium formate and 0.125% formic acid were added to both solvents. For chromatographic separation of polar and semi-polar metabolites, the resuspended samples were injected (1 µl) onto a Waters Acquity UPLC BEH Amide column (150 mm× 2.1 mm; 1.7 µm) with an additional Waters Acquity VanGuard BEH Amide pre-column (5 mm× 2.1 mm; 1.7 µm) held at 45°C. The separation of polar metabolites was achieved with a flow rate of 0.4 mL/min under the following gradient: maintained at 100% B (0-2 min), B changed linearly from 100% to 70% (2-7.7 min), B decreased to 40% (7.7-9.5 min), B decreased to 30% (9.5-10.25 min), B increased to 100% (10.25-12.75 min) and maintained at 100% B (12.75-16.75 min) for equilibration. Spectral data was collected with a scan range of 50–1200 m/z. MS/MS fragmentation was obtained using data-dependent acquisition (DDA) for the top five most abundant ions from each MS1 scan. The same Orbitrap source parameters were the same as lipidomics analysis. Normalized collision energies of 15, 30, and 45 eV were used as the collision energy.

### 2.11. LC-MS/MS and GC-MS data processing using MS-DIAL

Data were reported as quantitative ion peak heights. Data processing for data from each of the three analytical platforms (RPLC-MS/MS, HILIC-MS/MS, and GC-MS) was performed using the open-source software MS-DIAL (version 4.8) and included peak detection, deconvolution, feature alignment, gap filling, deisotoping, adduct identification, accurate mass/retention time (m/z-RT) library matching, and MS/MS library matching [1]. Features were annotated using defined confidence levels [2].For lipidomics, experimental MS/MS spectra were matched to reference MS/MS spectra from the LipidBlast library [3]. For HILIC data sets, experimental tandem MS spectra were matched to library spectra from the Mass Bank of North America (MoNA) (http://massbank.us) and NIST 17 MS/MS spectral library to annotate the features [4]. GC-MS spectra were matched against public mass spectral libraries (Fiehn library, In-house library, MassBank Japan, GOLM DB, GNPS, and HMDB). The GC-MS spectra were annotated to metabolite names by using two orthogonal parameters, such as high mass spectral similarity and retention indices (RI). Retention indices were calculated using an internal alkane standard. For GC-MS data, the signal intensity for each feature was normalized to the sum of intensities of all identified metabolites (mTIC normalization) to scale each sample before proceeding with statistical analysis [5]. For lipidomics, HILIC and GCMS in-house accurate mass and retention time m/z-RT libraries (created with authentic standards under identical chromatography conditions as it was used in this study) were used for metabolite identification. Duplicate peaks, isotopes, and adducts were removed using Mass Spectral Feature List Optimizer (MS-FLO) [6]. Peak height was used for quantification, and missing values were replaced with the minimum feature intensity divided by 10. Blank subtraction was carried out by removing features that had a maximum sample intensity/average blank intensity ratio of less than 3.
